# Entropy change due to selection in replicator and recombinator systems with randomly chosen interaction matrix values

**DOI:** 10.1101/2022.04.05.487241

**Authors:** W. A. Tiefenbrunner

## Abstract

Selection can be described as a process of information accumulation^21^. The present work extends this for frequency-dependent selection. For recombination, it is shown that the total entropy change can be calculated by summing the contributions of the individual gene loci in this case as well.

For more complex replicator systems with and without recombination with randomly selected interaction matrix values (defining the fitness landscape), a detailed investigation is made of how the number of replicators (“genotypes”) surviving, the mean fitness, and the entropy change depend on the number of replicators initially present, on their initial frequencies, and on certain properties (symmetry: a_ij_=a_ji_ versus a_ij_≠a_ji_) of the interaction matrix. Survival of several to many replicators is the norm rather than the exception. The more survive, the smaller the entropy change due to selection, although this is no longer generally true for asymmetric interaction matrices. Cases deviating from the rule were studied, but their systematic classification is still lacking. Finally, how evolution alters replicator systems of greater complexity was analyzed. It is shown that different evolutionary algorithms can lead to an increase in mean fitness and/or an increase in the number of surviving replicators. In contrast to simple replicator systems, coexistence in more complex ones apparently results naturally as a consequence of selection and evolution. It results that the view that: “Evolution is based on a fierce competition between individuals and should therefore reward only selfish behavior”^16^ is merely an artifact that follows from the study of unrealistically simple systems.

## Introduction

In 1983, Schuster and Sigmund^19^ made the astonishing discovery that the mathematical description of self-organization in seemingly quite different fields is based on the same differential equation. Later they also gave it a name - replicator equation - and in the mentioned paper they called the mathematical framework “replicator dynamics”. A replicator is information that can be copied. This information, the message, must exert an influence on the frequency with which it is copied. This can happen directly or indirectly, in that the properties of the carriers of the information are determined by the message, and these properties in turn influence the probability of copying. The replicator equation (Nowak^15^ also calls it the equation of density-dependent selection) states:

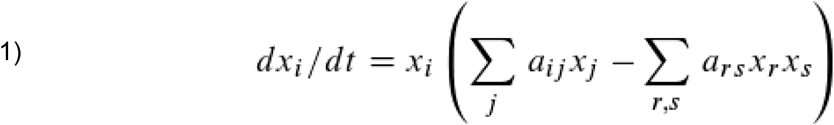

where i,j,r,s=1…n and for all a_ij_ holds: a_ij_≥0. Here x stands for the relative frequency of replicators and a for the interaction coefficient describing the interaction between two replicators. n is the number of different replicators. The continuous Eq. 1 corresponds to the difference equation:

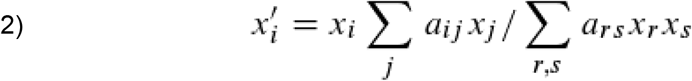

with x_i_’=x_i_+Δx_i_/Δt. Δt can be, for example, the duration of a generation, Δx_i_ the change in frequency in this time interval. The equation is such that from Σx_i_ =1 follows also Σx_i_’ =1. The authors mention four fields of research in which this formula plays a role, in each of which it was discovered independently.

1. In population genetics, it provides the selection equation^4^ (with symmetric interaction matrix), which describes the frequency of alleles in the gene pool. An allele is a replicator, whereas the genotype, when recombination occurs, is not (I refer to it here as a recombinator).
2. In mathematical ecology it represents as Lotka-Volterra equation^11, 24^ the relative numbers of individuals of species or populations. Individuals differ from each other, so strictly speaking they are not replicators, but the differences within the population are small enough to treat them as if they were in an idealized view.
3. As a rate equation in the flux reactor and to describe a hypercycle, the replicator equation is of great importance in the field of prebiotic evolution, modeling the concentrations of self-replicating, information-bearing macromolecules^2^.
4. Finally, in sociobiology, it appears as a game dynamic equation in the description of heritable or tradable conflict behavior^12-14^.

In 1983, Haken^5^ already pointed out that the equations describing the autocatalytic propagation of biomolecules in the original version of Eigen’s theory are identical to those dealing with the amplification of light waves or photons in his laser model^6-9^. Thus, also photons, resp. more generally: bosons, behave under certain circumstances like replicators (a realization, which actually already follows from a paper of Einstein 1917^3^), with which the area, in which the replicator dynamics is of relevance, extends up to quantum mechanics and thus pervades the entire natural science from physics to behavioral research.

Eigen^2^ discovered an essential property of replicators in 1971: Even if the copying process is erroneous, i.e. subject to statistical noise, the replicator information can in principle remain unchanged for an infinitely long time under certain conditions due to the propagation and corresponding selection resulting from it. Realistic are at least long periods of time, as demonstrated e.g. by Homer’s Illiad and Odyssey, which have existed for at least 2600 years. This example also shows very clearly that the actual replicator is the information and not its carrier. The Illiad occurs on parchment and paper, but also as the memory content of hard disks and other storage media, making it one of the most successful artificial replicators. However, success is not relevant for the definition: this article is also a replicator in principle. The conditions for information to be preserved even if copying errors occur were formulated by Eigen^2^ as the “error threshold”. It can also be understood as a complexity threshold, since it describes how much information is preservable, given copying error on the one hand and copying success on the other.

Another contribution to replicator dynamics was made by Tiefenbrunner 1995^21^ by showing that selection based on competition can also be described in a completely different way, namely as a process of information acquisition. I.e., to a selection step described by the discrete equation

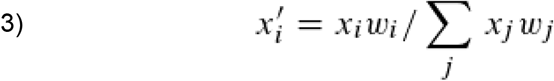

an information change ΔH per generation (or time step) can be determined in the form:

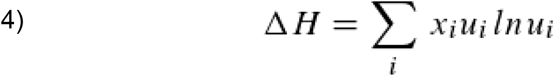

with

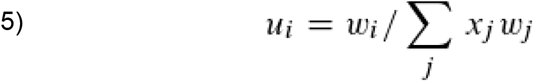

Here, w_i_ is the fitness (the degree of adaptation to the environment) of genotype X_i_, which is assumed to be constant. Σx_i_w_i_ is therefore also called “mean fitness”, where x_i_ is of course the relative frequency of X_i_. u_i_ is the “distinctness” of X_i_ and again i,j=1…n. Because fitness is very difficult or impossible to determine in practice, an alternative formulation of the change in information content due to selection is often more appropriate for practical application^22^:

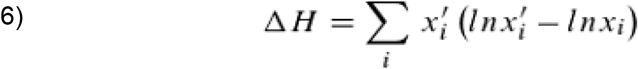

In the same work, it was shown for recombinators that if there are j gene loci (here j=1…m, so j does not index the replicators or recombinators as i does) and recombination occurs without linkage:

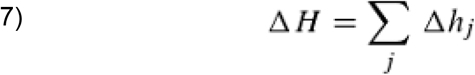

Δh_j_ is the information change per gene locus:

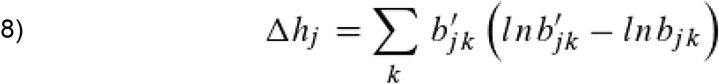

Here B_jk_ is the kth allele of the jth gene locus and b_jk_ its relative frequency in the population (k is defined by i and j, as described in Supplement 1). This means that the total information change per generation by selection can be determined from the contributions of each gene locus simply by summation (as long as it is the own genotype that determines the phenotype and not, for example, the maternal one^22^). This is quite remarkable and probably the most fundamental property of a recombinator system (i.e., a system in which, in addition to selection, the laws of recombination are valid, occurring without linkage). For the mean fitness ω= Σx_i_w_i_, which like the information change ΔH characterizes the entire recombinator system, this is only true if the overall fitness behaves additively, i.e., if the fitness of the genotype is equal to the sum of the fitness values of the alleles of the individual gene loci composing it. As an example this is shown for two loci in Supplement 2.

Do Eqs. 7 and 8 apply only to the classical selection equation (Eq. 5) or also to Eq. 2, which describes density-dependent selection? One gets from Eq. 5 to Eq. 2 by defining:

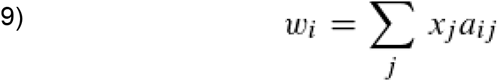

i,j=1…n, where n is the number of genotypes. Now w_i_ does not occur at all in Eq. 6-8, nor does it occur in the original derivation of these equations. Thus, one can actually make a substitution in the form Eq. 9 without affecting the correctness of Eq. 7. Fig. 1 shows this exemplarily for four gene loci with two alleles each in a model calculation based on the given equations.

**Fig. 1:**
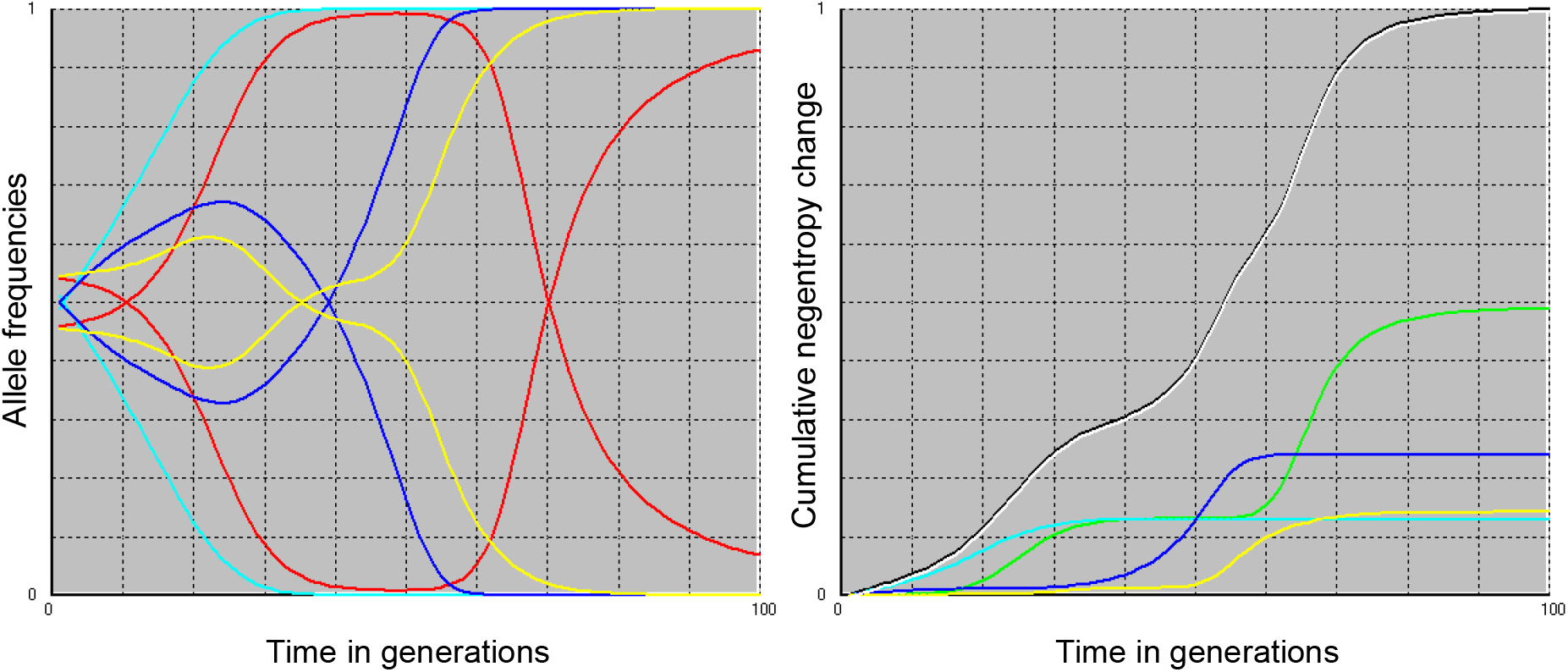
Left figure: Selection within 100 generations. Involved are sixteen genotypes (recombinators), which correspond to all possible combinations of 4 gene loci to 2 alleles each. In different colors the allele frequencies of different loci are shown, in the same colors the two alleles of one gene locus whose frequencies complement each other to 1 (hence the symmetry). The frequency change of the eight alleles is presented. Right figure: cumulative negentropy change of the different loci during the selection process (in colors), sum of these values over all four loci (white) and cumulative negentropy change obtained when genotype frequency is used as a basis for calculation (black). The white and black curves are drawn slightly offset from each other, as they would otherwise completely overlap.

More follows later regarding the choice of initial frequencies of genotypes and values of the interaction matrix.

An interesting question is the meaning of ΔH, but also ω in the context of replicator dynamics. As long as w_i_ was seen as a constant, ΔH was very easy to interpret as a change in information content due to the selection process. But if w_i_ is described by Eq. 9, what is the meaning of ΔH? In any case, until this question is resolved, it is no longer meaningful to speak of information content, and we therefore use only the term “negentropy.”

By the way, the same question about the meaning actually arises for ω, which was also formulated for the selection process with constant w_i_ and called mean fitness. By tradition, we retain the term here (although the author would prefer the term “intensity”).

In any case, ΔH, unlike Δω, is not symmetric with respect to the time axis. That is

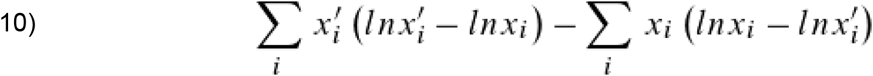

need not be zero, which is especially important for cyclic processes.

### Replicator systems with random interaction matrix values

A replicator system consists of multiple replicators interacting with each other. For example, one can imagine short, t-RNA-like molecules that simultaneously carry information but are also ribozymes, carriers of catalytic properties. In this way, one escapes the vexing problem of constantly having to distinguish between genotype and phenotype; they are both at the same time.

A replicator system consists of X_i_ (i=1…n) different replicators and is defined by the number n of replicators present before the start of selection, their frequency at the beginning x_i_(t=0) and the constants a_ij_ of the interaction matrix, i.e. the mutual influence of their reproduction or survival probability. The current state of the replicator system can be characterized by the mean fitness (intensity) ω, the selection process by the change in mean fitness Δω, the change in negentropy ΔH and, of course, by the change in frequencies x_i_ and, consequently, by the extinction of some replicators.

A great many papers have been written on the topics of cooperation or coexistence of replicators (e.g., refs. 1, 16, 17, 19). For n=2, coexistence occurs only when each replicator supports the other more than itself. For n>2, more complex scenarios are possible, in particular, one replicator can no longer easily behave “selfishly” toward another because that can indirectly affect it back through other replicators.

The possible results of selection are therefore very diverse, for example, only one replicator can survive (“survival of the fittest”) or several, which then promote each other, thus actually representing a “synreplicator” consisting of several replicators, whose complexity is given by the number of surviving, interacting replicators. Cyclic solutions are also possible (“scissors, stone, paper” and others) and finally chaos^10,15^.

The following model calculation is based on the question how selection affects a replicator system with initially n replicators (n=4, 8, 16, 32), where the interaction values a_ij_ (i,j=1…n) are chosen randomly in the range 0<a_ij_≤1. However, in a variant of the experiment, a_ij_=a_ji_ should be required, i.e., the interaction matrix should be symmetric (the notion of symmetry refers exclusively to the property just defined, i.e., even an asymmetric interaction matrix is a square matrix).

As for the initial frequencies of the replicators, let us also examine two situations: In one case we want to assume that for all i holds: x_i_=1/n (hence Σx_i_=1), i.e., initially all replicators are equally frequent. In the other, we want to specify that the frequencies x_i_ are random at the beginning, but of course the condition Σx_i_=1 must hold in this case as well. The selection is to be done according to Eq. 2. This fully describes the replicator system and we can now consider what questions to ask the model. First, we might be interested in how many different replicators are left after, say, 1000 generations, or how many replicators make up the “surviving” synreplicator (what its complexity k is) and how the frequency of k (which we will denote as c_k_) depends on the number of replicators at the start of the experiment, on the symmetry (or asymmetry) of the interaction matrix, and on the initial frequencies x_i_. Fig. 2 shows the result, where the experiment was repeated 10000 times and after 1000 generations in which selection had taken place, it was determined how often (corresponding to c_k_) a certain synreplicator complexity k (k=1…n) had been reached (i.e. how many replicators survived and how often this was the case).

**Fig. 2:**
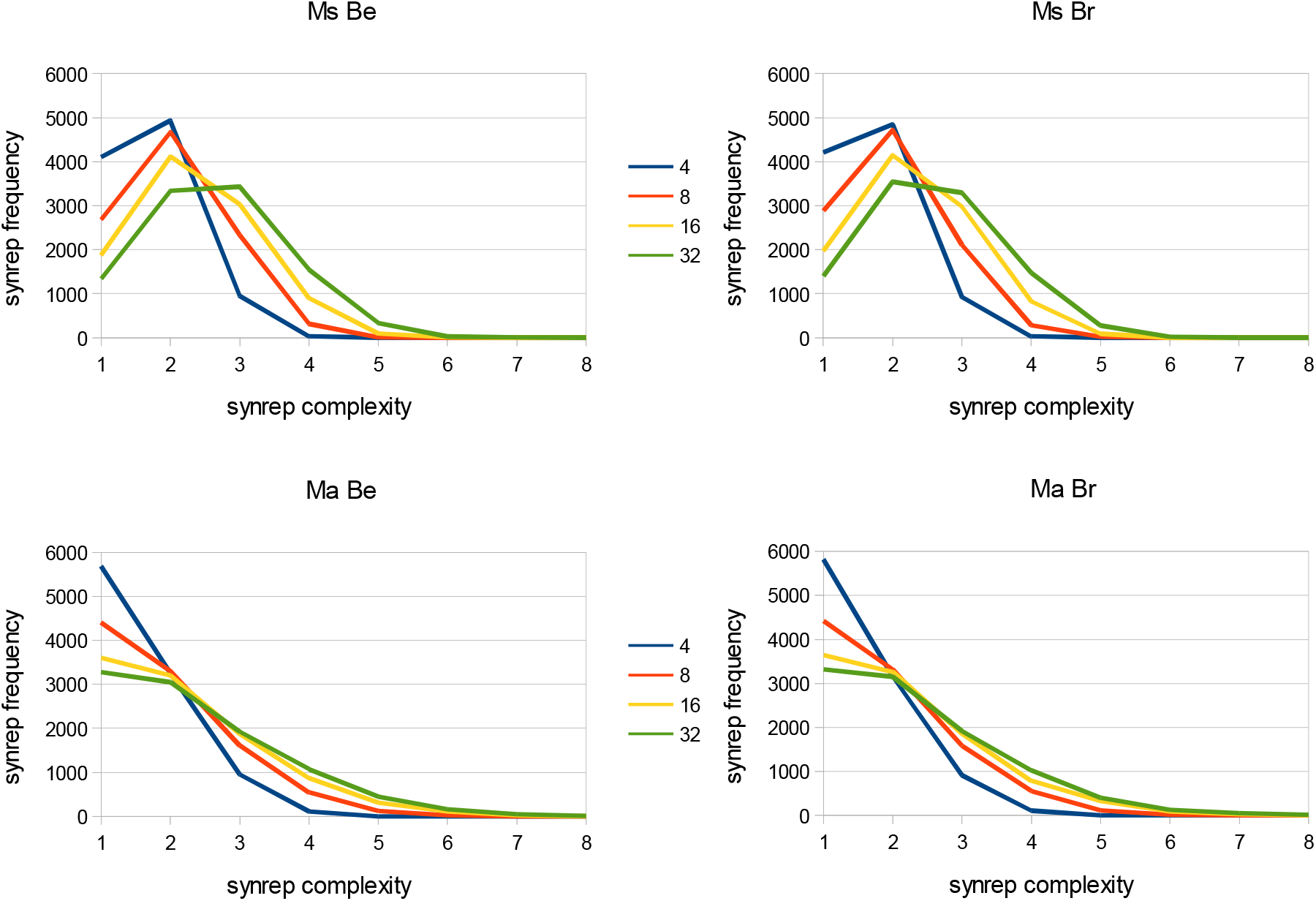
Number of surviving replicators (synrep complexity) k and their frequency (synrep frequency) c_k_ after 1000 generations in which the replicator system (with n=4, 8, 16, 32 replicators at the beginning; curves with different color) has been subjected to selection according to Eq. 2. The experiment was repeated 10000 times. See text for further explanation. Ms: symmetric interaction matrix a_ij_=a_ji_ ; Ma: asymmetric interaction matrix; Be: initial frequencies x_i_ of the replicators is equal to 1/n for all; Br: random initial frequencies.

Fig. 2 distinguishes four cases resulting from the combination of symmetric or asymmetric interaction matrix and equal (x_i_=1/n) or random initial frequency of replicators.

In the case of symmetric matrix, more than one replicator survives in over 50% of the repetitions (k>1), sometimes up to seven. As expected, the frequency c_k_ with which one observes a given synreplicator complexity k depends only to a very small extent on whether the initial frequencies x_i_ of the replicators were chosen randomly or not. In contrast, the number n of replicators initially present is very important. The larger it is, the less often the survival of only one or two replicators is observed; with an initial n=32 different replicators, a survival of a single replicator is only observed in about 13% of the replicates, a synreplicator complexity of k=2 just in 33%. In contrast, for a synreplicator complexity of three or more (k≥3), the number c_k_ grows with the initial replicators, which is in part simply a consequence of the fact that in a system with more replicators initially, there are naturally more possible synreplicator complexity levels (namely, n). In the context of the simulation described here, the maximum number of surviving replicators was k=5 for replicator systems of n=8 components, k=6 for n=16 components, and finally k=7 for n=32 initially present replicators. Thus, it increases with the size of the replicator system, albeit nonlinearly. Overall, synreplicators of complexity two or three are observed most frequently.

That only one replicator survives the selection process occurs more frequently with asymmetric interaction matrix, with initial n=4 the frequency is even clearly above 50% (c_1_=57% with equal initial frequency of the replicators and c_1_=58% with random initial frequency). Already at n=8 it is below 50%, but still remains above 30% even at n=32. Whether the initial frequencies x_i_ of the replicators are random or equal plays only a very minor role for the distribution of the synreplicator complexity. A complexity k=2 is reached almost independently of n with a frequency of c_2_=30% (n=32) to c_2_=33% (n=4). Only when k>2 does c_k_ again depend significantly on n: the larger n, the greater the frequency c_k_ of a given synreplicator complexity level k. Comparing c_k_ of symmetric and asymmetric matrix, it is noticeable that up to k=4 the synreplicator frequency is significantly higher for symmetric matrix (c_4_≈15% versus c_4_≈10% to 11%), while from k=5 the ratio reverses (e.g., c_5_≈3% for symmetric versus c_5_≈4% to 5% for asymmetric interaction matrix). Accordingly, larger k also occur with asymmetric matrix (not shown in Fig. 2). k=9 and even k=10 have been observed for n=32 when the interaction matrix was asymmetric, and k=8 is even relatively frequent (c_8_=0.14%, i.e., as many as 14 observations for 10000 repetitions).

The next question we ask the model is about the mean fitness ω_k_ after 1000 generations of selection: how does it change with n (the number of replicators initially present) and k (the synreplicator complexity), and what is the influence of matrix (a)symmetry and initial frequencies (Fig. 3)? For random interaction values 0<a_ij_≤1, we can expect the intensity ω to be close to 0.5 before selection. Values lower than this will thus be unlikely to be observed after selection has occurred. Values greater than one are also not possible. The highest mean fitness after selection is observed at n=32 and k=1, but it is hardly even higher than at n=16. In general, the larger the number n of replicators initially present and the smaller the synreplicator complexity k, the larger is ω_k_. Symmetry also plays a major role. Smaller values of ω are obtained for asymmetric matrix. For n=k=4 it is even true that ω_4_ is not noticeably higher after selection than before. Whether or not the initial frequencies of the replicators are chosen randomly is hardly important.

**Fig. 3:**
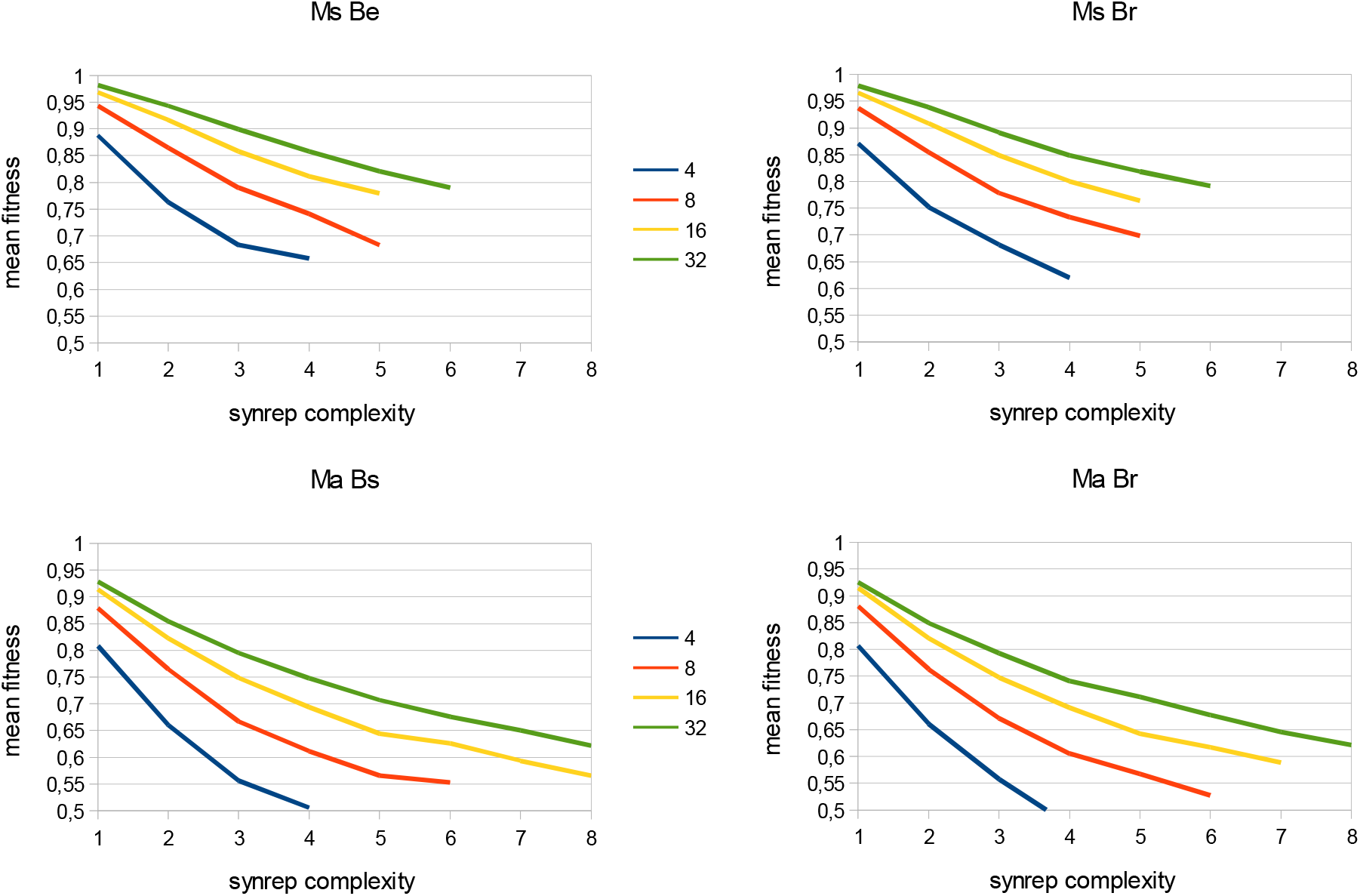
Number of surviving replicators (synrep complexity) k and the mean fitness ω_k_ after 1000 generations in which the replicator system (with n=4, 8, 16, 32 replicators at the beginning; curves with different color) has been subjected to selection according to Eq. 2. The experiment was repeated 10000 times. See text for further explanation. Ms: symmetric interaction matrix a_ij_=a_ji_ ; Ma: asymmetric interaction matrix; Be: initial frequencies x_i_ of the replicators is equal to 1/n for all; Br: random initial frequencies.

Because of the close relationship between mean fitness ω_k_ and change in negentropy ΔH_k_ (eqs. 4 and 5), one might think that the latter behaves essentially similarly to the former in the model. Fig. 4 shows that this is certainly true for the symmetric interaction matrix. The relationship between cumulative entropy change and k might even be linear for a given n, and ΔH decreases with n and k. In the case of the asymmetric interaction matrix, however, this is not the case at all. Only up to k=2 does the cumulative negentropy ΔH (always referred to 1000 generations of selection) decrease, but then it increases again, and the magnitude of this increase, moreover, apparently depends on the initial frequencies x_i_ (compare Fig. 4 Ma Be with Ma Br). Moreover, n also plays a role: from k=4, the smaller n is, the steeper the increase in cumulative entropy. We will look for the reason for this peculiar behavior in a while, but before that we will complete this investigation for recombinator systems.

**Fig. 4:**
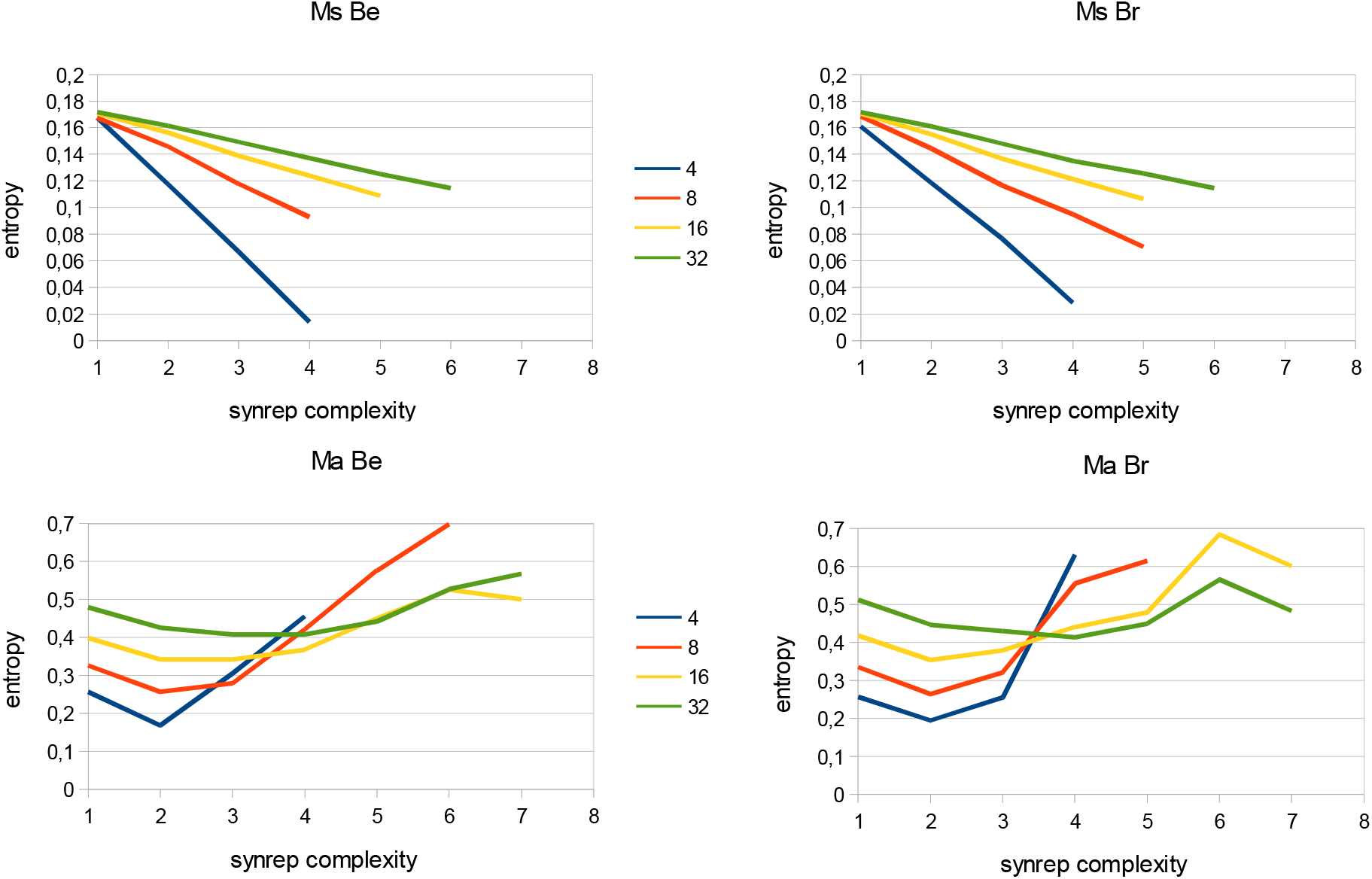
Number of surviving replicators (synrep complexity) k and cumulative negentropy change (simplified as “entropy”) ΔH_k_ over 1000 generations in which the replicator system (with n=4, 8, 16, 32 replicators at the beginning; curves with different color) has been subjected to selection according to Eq. 2. The experiment was repeated 10000 times. See text for further explanation. Ms: symmetric interaction matrix a_ij_=a_ji_ ; Ma: asymmetric interaction matrix; Be: initial frequencies x_i_ of the replicators is equal to 1/n for all; Br: random initial frequencies.

### Recombinator systems with random interaction matrix values

In addition to pure replicator systems, there are also conceivable systems in which recombination (always without linkage) is allowed (“recombinator systems”). In this case, allele number l and number of gene loci m are added as factors necessary for a complete description of the system. For the number of “genotypes”, recombinators, n=l^m^ holds. If we work with l=2 and m=2 to 5, we get n=4, 8, 16, 32 (this explains why we started from these initial numbers for the model calculations even for replicator systems; it is for comparison).

Note that in recombinator systems, it is the alleles that die out. Therefore, in the context of the model, the number of recombinators k surviving after 1000 generations of selection can only take the values 1, 2, 4, 8, 16, and 32, which is a difference from replicator systems that should be noted and has been considered in Figs. 5 to 7 below. The model calculations were repeated 10000 times and each time the relevant parameters k, ω and ΔH were determined, which finally allowed to plot the distributions c_k_ (Fig. 5), ω_k_ (Fig. 6) and ΔH_k_ (Fig. 7).

**Fig. 5:**
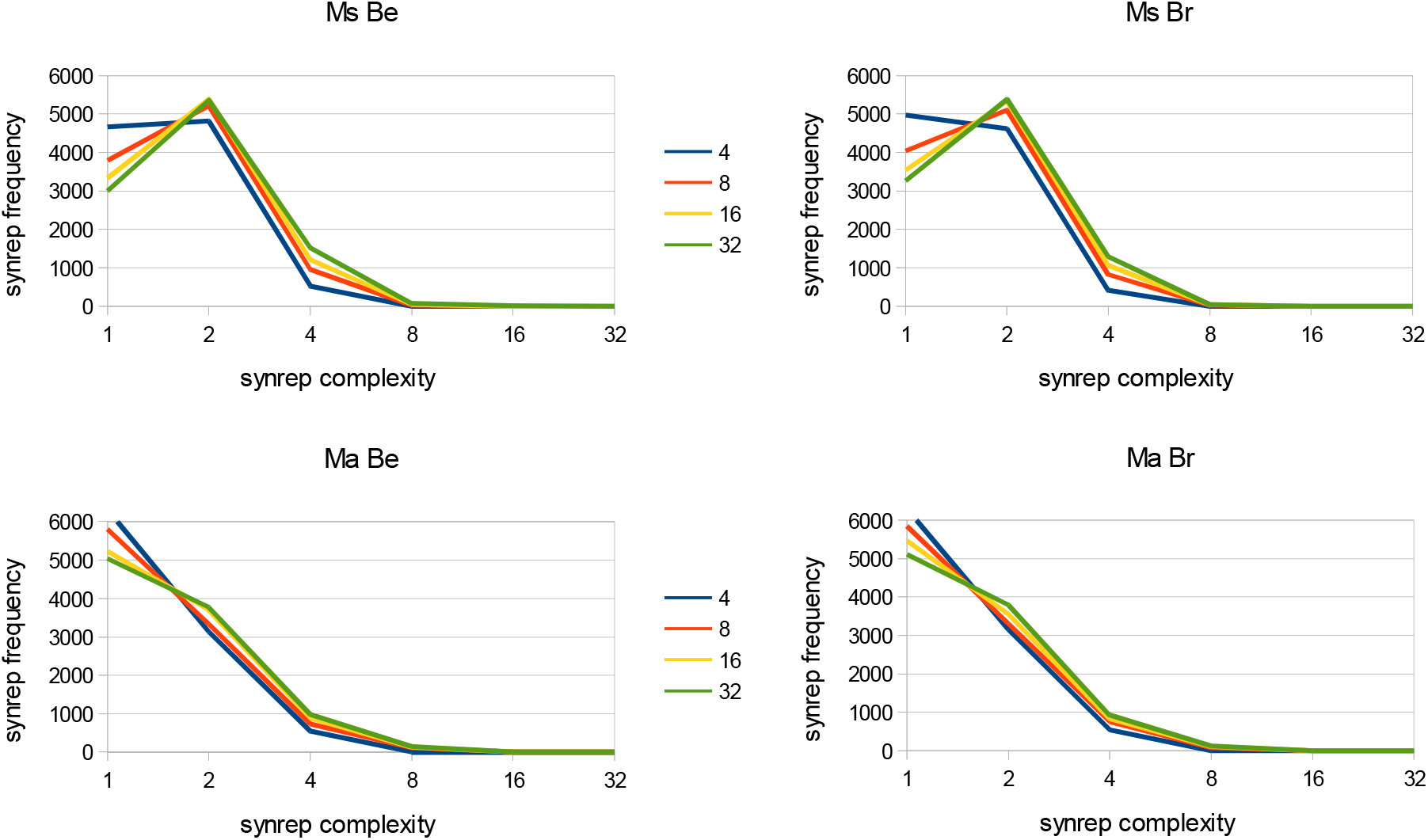
Number of surviving recombinants (synrep complexity) k and their frequencies (synrep frequencies) c_k_ after 1000 generations in which the recombinator system (with n=l^m^=2^2^, 2^3^, 2^4^, 2^5^=4, 8, 16, 32 recombinators at the beginning; curves with different color) has been subjected to selection according to Eq. 2. The experiment was repeated 10000 times. See text for further explanation. Ms: symmetric interaction matrix a_ij_=a_ji_ ; Ma: asymmetric interaction matrix; Be: initial frequencies x_i_ of the replicators is equal to 1/n for all; Br: random initial frequencies.

**Fig. 6:**
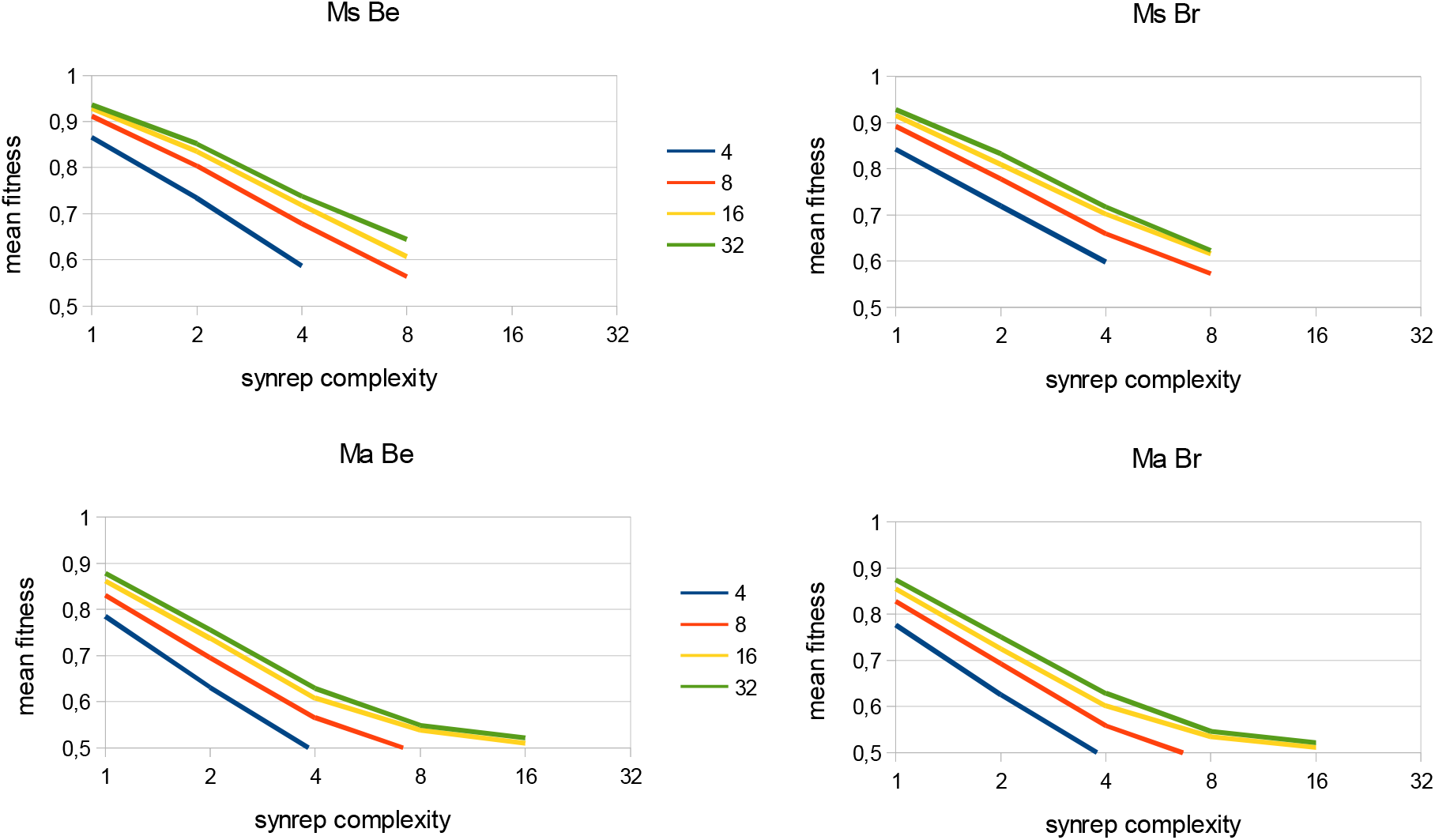
Number of surviving recombinators (synrep complexity) k and mean fitness ω_k_ after 1000 generations in which the recombinator system (with n=l^m^=4, 8, 16, 32 recombinators at the beginning; curves with different color) has been subjected to selection according to Eq. 2. The experiment was repeated 10000 times. See text for further explanation. Ms: symmetric interaction matrix a_ij_=a_ji_ ; Ma: asymmetric interaction matrix; Be: initial frequencies x_i_ of the recombinators is equal to 1/n for all; Br: random initial frequencies.

**Fig. 7:**
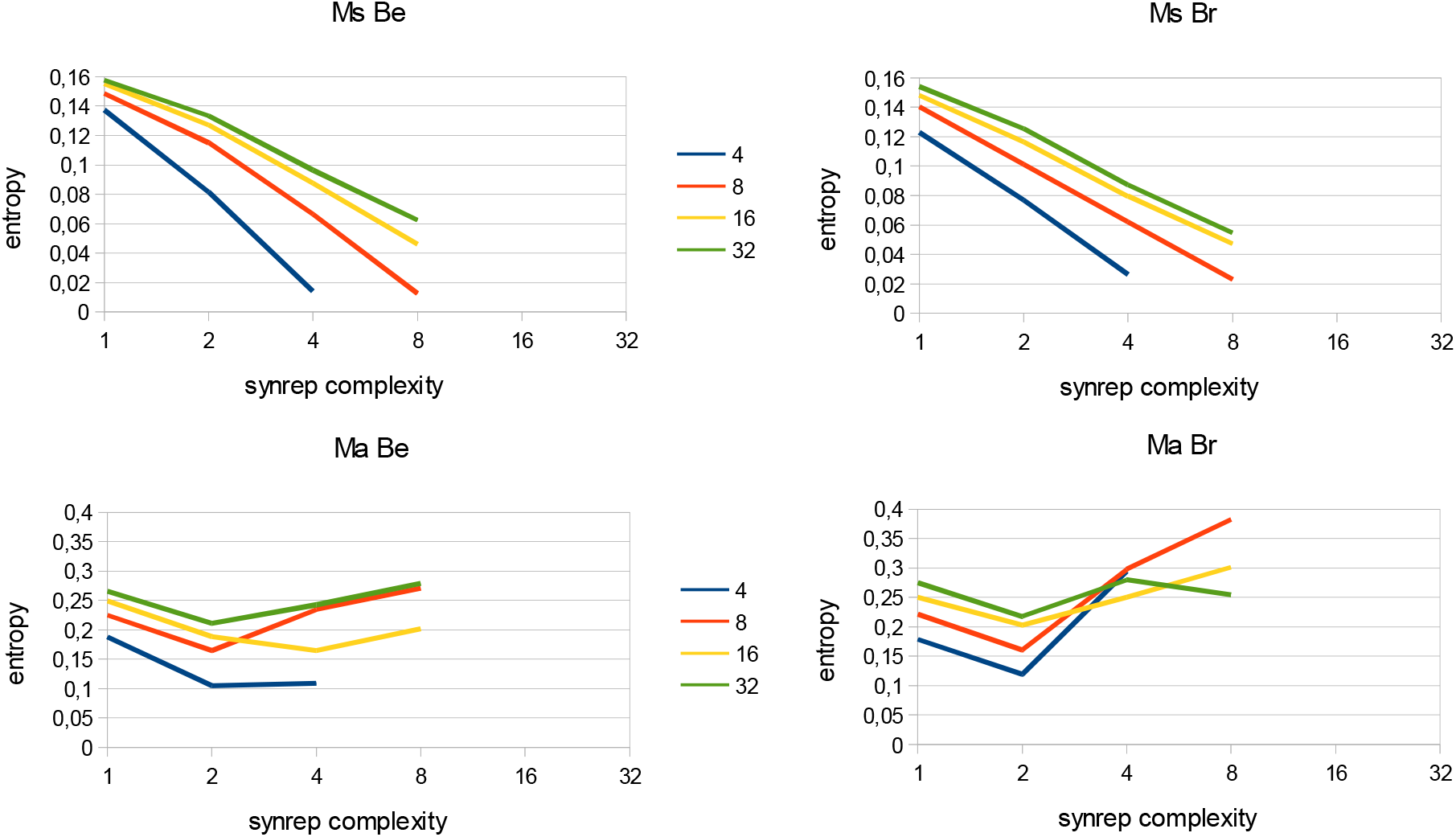
Number of surviving recombinators (synrep complexity) k and cumulative negentropy change (simplified as “entropy”) ΔH_k_ over 1000 generations in which the recombinator system (with n=l^m^=4, 8, 16, 32 recombinators at the beginning; curves with different color) has been subjected to selection according to Eq. 2. The experiment was repeated 10000 times. See text for further explanation. Ms: symmetric interaction matrix a_ij_=a_ji_ ; Ma: asymmetric interaction matrix; Be: initial frequencies x_i_ of the recombinators is equal to 1/n for all; Br: random initial frequencies.

Similar to replicator systems, in the simulation with recombinator systems with **symmetric matrix** more than one recombinator (k>1) survived in more than 50% of the repetitions, rarely (e.g. 64 out of 10000 with n=32) up to eight, in one case even k=16. It doesn’t matter much whether the initial frequencies x_i_ of the recombinators are chosen randomly or not, but with the same frequency at the beginning k=8 was found more often. The number n of initial recombinators is not as important as the number of replicators in replicator systems and matters most for k=1. The larger n, the less often only one recombinator survives. A synreplicator complexity of k=2 is observed in about 50% of the repetitions. For a synreplicator complexity k of three or more, the c_k_ grows only slightly with the number of initial recombinators.

There is agreement between replicator and recombinator systems in that it is more common for only one “genotype” (in this case, a recombinator) to survive the selection process when the interaction **matrix** is **asymmetric**, i.e., only one allele at each gene locus survives.

At n=4 the frequency c_1_ is 63% and even at n=32 it is still more than 50%, which is a clear difference to replicator systems. Whether the initial frequencies x_i_ of the replicators are random or equal has little effect on what happens. The importance of n is less than in replicator systems, but more “genotypes” survive even in asymmetric matrix (k=16 was observed in 6 repetitions out of 10000 with initial n=16 or more recombinators, with equal initial frequencies it were even 12 repetitions), also more than in symmetric matrix in recombinator systems. From a complexity of k>2, the difference between symmetric and asymmetric matrix is relatively small compared to replicator systems.

How the mean fitness ω_k_ changes after 1000 generations of selection as a function of n (the number of replicators initially present), k (the synreplicator complexity), matrix (a)symmetry, and initial frequencies is shown in Fig. 6. For random interaction values 0<a_ij_≤1, the intensity ω is generally close to 0.5 at the beginning and smaller values are not observed after selection has occurred. Values larger than one are impossible. Thus, as in replicator systems, the highest mean fitness ω_1_ after selection is found at n=32 and k=1, differing little from that at n=16. ω_k_ is larger the larger the number n of recombinators initially present and the smaller the synreplicator complexity k. The symmetry of the interaction matrix is essential and has very similar effects as in replicator systems. Slightly smaller values of ω are generally obtained for asymmetric matrix. For n=k=4, as well as n=k=8, ω_4_ and ω_8_, respectively, are not higher after selection than before. Whether the initial frequencies of the replicators are chosen randomly or not turns out to be unimportant.

Fig. 7 shows that the mean fitness ω and the change in negentropy ΔH behave similarly for symmetric interaction matrix in the model, as expected from Eq. 4 and Eq. 5. In this respect, replicator and recombinator systems agree, but differ in that in recombinator systems the relationship between cumulative entropy change ΔH_k_ and k is certainly nonlinear for given n (note that the x-axis showing the distances from k is nonlinear in Fig. 7).

For asymmetric interaction matrix, the curve representing ΔH_k_ takes trough shape, much like in replicator systems as well. That is, only up to k=2 does the cumulative negentropy ΔH decrease (in each case after 1000 generations of selection), but then it increases again, and the magnitude of this increase also appears to depend on the initial frequencies x_i_ (compare Fig. 7 Ma Be with Ma Br). While for symmetric matrix ΔH decreases with n and k, the situation is apparently not so simple for asymmetric interaction matrix from k=4 onwards.

As we have seen, qualitatively the results for replicators and recombinators are mostly in agreement.

Have we asked all the important questions that can be asked about selection in a replicator or recombinator system with random interaction values a_ij_? The answer is no. This is because there is often more than one possible final state depending on the initial values x_i_ for a given interaction matrix. The question is then, of course, how many different “solutions” occur, what synreplicator complexity k they exhibit, and with what frequency they are distributed over k. As interesting as this question is, it does not contribute anything new to learn about entropy change and is therefore excluded from this paper.

### Increase in cumulative negentropy with asymmetric interaction matrix and k≥3

Looking for model calculation examples where the negentropy increases unexpectedly fast, one will find those where no stable, stationary equilibrium between the frequencies of the replicators (recombinators) is established during selection or where this is the case only with a delay. Chaotic and cyclic behavior of the replicator or recombinator system, which is possible only when the interaction matrix is asymmetric, is thus the cause of the unusual increase in negentropy – and thus Eq. 10, which states that ΔH, unlike Δω, is not symmetric with respect to the time axis. Examples are provided in Fig. 8, although it is also apparent that not every oscillatory or damped oscillatory system leads to a continuous or excessive increase in negentropy. A great deal of research is undoubtedly still needed to achieve a classification of replicator systems with complex behavior, and negentropy is obviously a good pivot from which to approach this endeavor.

**Fig. 8:**
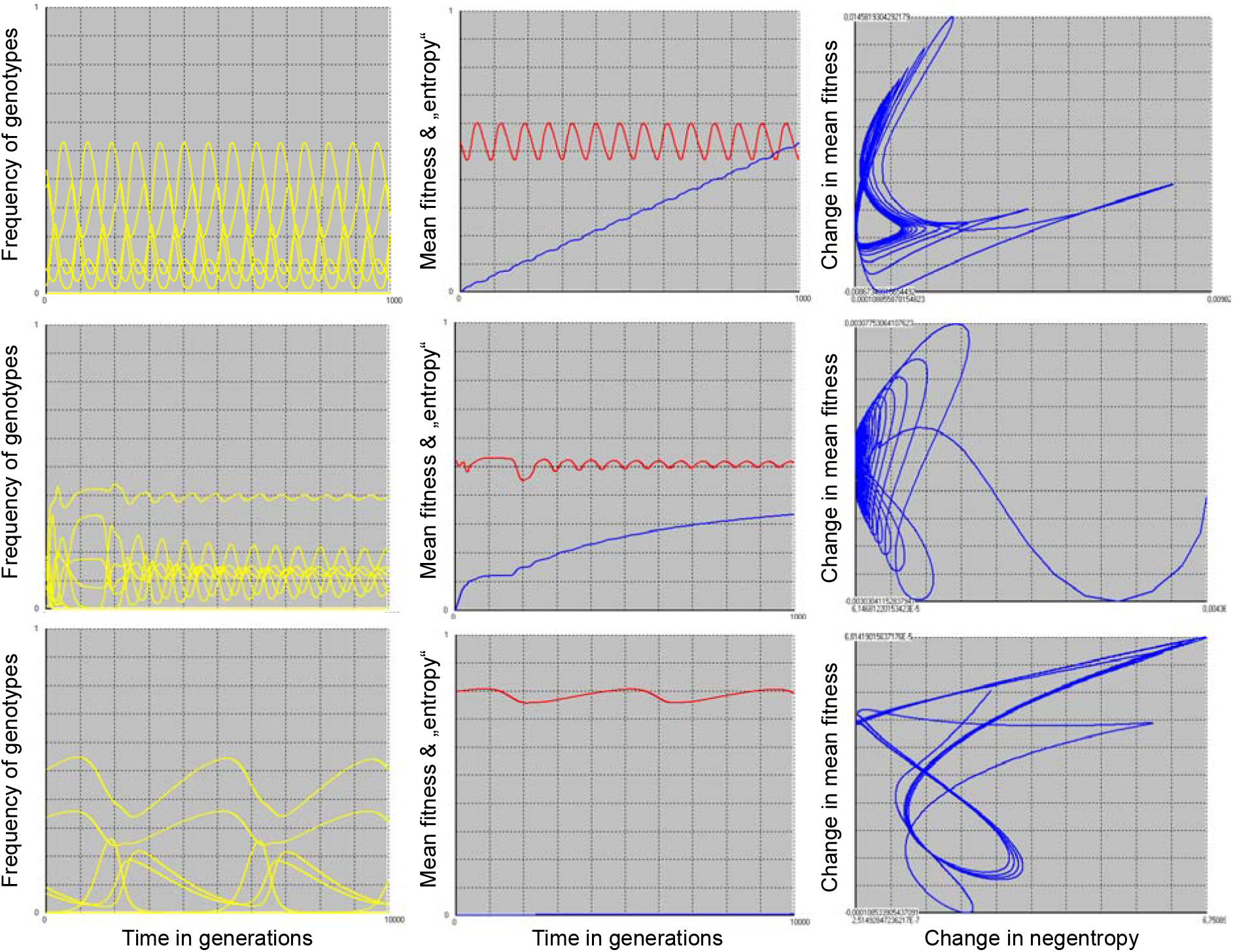
Replicator systems (n=10) with cyclic behavior. In columns: Left: relative frequency of surviving replicators versus time in generations. Middle: red: mean fitness; blue: cumulative change in negentropy (“entropy”) versus time in generations. Right: change in mean fitness versus change in negenetropy. Shown in rows are three different systems (top: continuously oscillating system with approximately constant negentropy rate and therefore linear increase in cumulative negentropy; middle: damped oscillating system whose negentropy rate will tend to zero but is still positive; bottom: continuously oscillating system with negentropy rate equal to zero).

Fig. 8 shows three examples of replicator systems with n=10 which do not reach a steady state or only with a strong delay. The top row presents cyclic behavior with large cumulative change in negentropy (blue curve in the middle column) with mean fitness also oscillating (red curve). The oscillation is also evident in the plot in the right column, where negentropy rate (x-axis) is plotted against the rate of mean fitness (y-axis). The attractor is a closed curve, with a negentropy rate of zero reached only once in the cycle (where the curve touches the y-axis).

The example in the central row presents a damped oscillation, with initially large change in negentropy. The right figure shows that the system tends to a negentropy rate of zero.

The bottom row presents an oscillating system where the negentropy does not change after the initial phase. In the right figure, the beginning of this process is shown, while in the other two, an advanced stage is already presented to clearly demonstrate the cyclic nature.

It should also be noted that the negentropy cannot increase in closed systems. From a physical point of view, the systems we study here are incompletely described. This is necessary because of their complexity.

### Evolution in replicator or recombinator systems with randomly chosen interaction matrix values

In the present model, each replicator is defined by its interactions with the other replicators and with itself, and by its frequency. The frequency can change by selection, the values of the interaction matrix have been assumed invariant so far. If one wants to allow evolution in the replicator system, one must also allow that the interaction matrix values become variable. Already up to now replicators could die out by selection. Now, new ones can also be added. We begin the simulation with randomly chosen relative initial frequencies for the n replicators, where, of course, the sum of all replicators equals one: If z_i_ is a random number 0<z_i_≤1, then xi=z_i_/Σz_j_. The sum runs from 1 to n. Also, the values of the interaction matrix are random numbers 0<a_rs_≤1 (and therefore the matrix is almost surely asymmetric with respect to the values). Selection is then performed for 1000 generations according to Eq. 2. The result is documented. For example, in Fig. 6 (yellow dots) one sees the number of surviving replicators (the synreplicator complexity) plotted against the mean fitness. A replicator is considered extinct when its abundance falls below a single individual, where we assume 1000 individuals total population size (so if x_i_<0.001). So far, everything runs as usual.

Now the reset to the initially selected replicator frequencies takes place, since a comparison should be possible. Subsequently, a certain process is repeated 1000 n^2^ times. This process consists of the alternating sequence of selection and mutation. Selection runs according to the described scheme. For mutation, different alternative algorithms were created and also combined with each other.

1. Mutation type 1 (“replace matrix value”) **replaces** a randomly chosen matrix value a_rs_ with a random number z (0<z≤1). If the replicator to which this value belongs is already extinct, a new initial frequency 1/n is set to give it a chance. When this mutation type is performed, there is a cross in the column “matrix” and the line “replace” in the legend to the individual figures of Fig. 9 (table in the lower right corner).

**Fig. 9:**
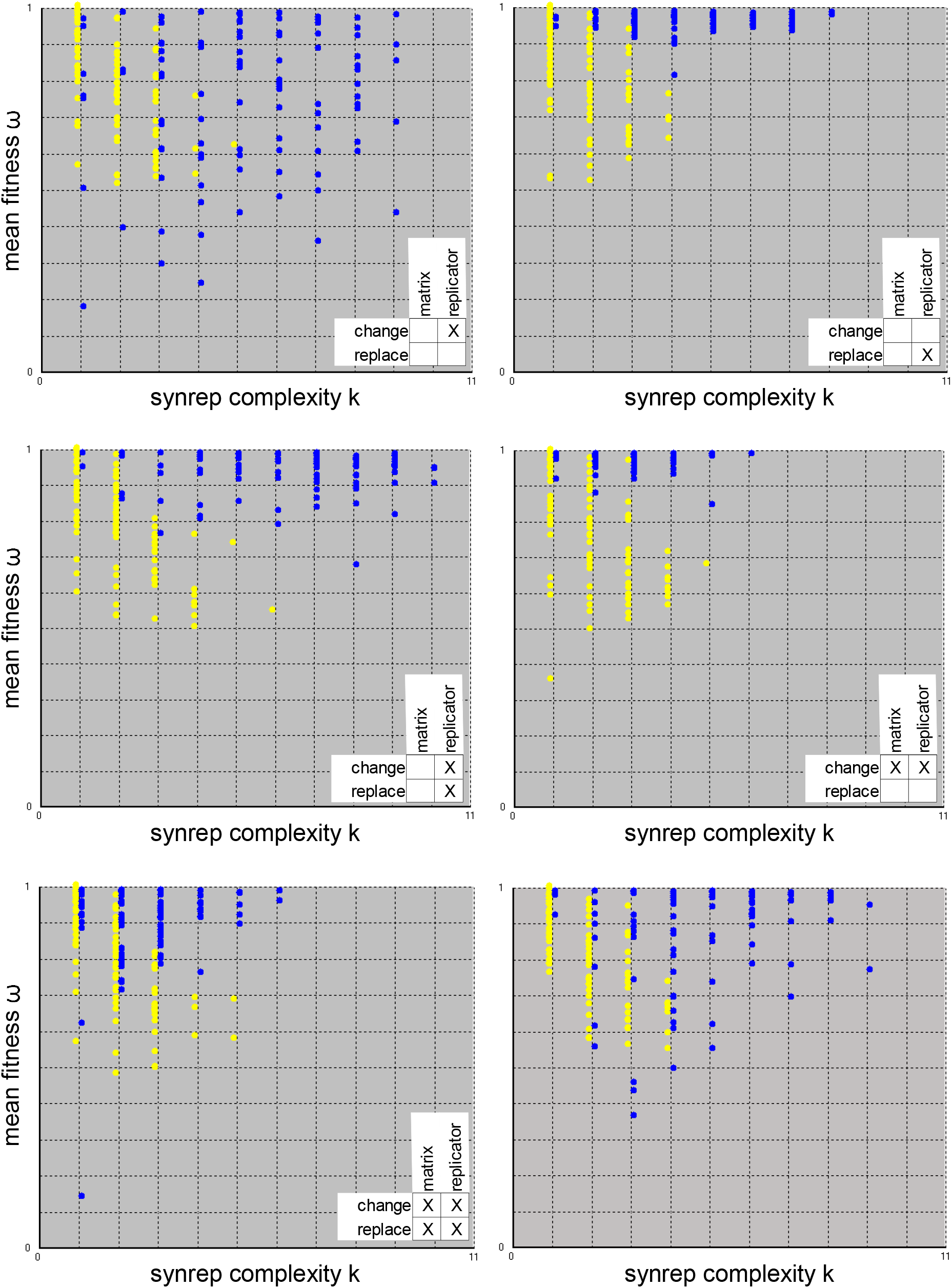
Comparison of replicator systems (n=10) with randomly chosen interaction matrix values after just selection (yellow dots) and after evolution (blue dots), i.e. selection and mutation. The insert at the bottom right of the individual figures indicates which mutation types were used. See text for further explanation.
2. Mutation type 2 (“change matrix value”) **mutates** a randomly chosen matrix value a_rs_: the value is increased or decreased by a maximum of 0.05 (a_rs_ +0.1z-0.05) or remains unchanged, if this operation would result in values below zero or above one for the changed a_rs_. If the replicator to which this value belongs is already extinct, the procedure is the same as for mutation type 1. If this mutation type is performed, there is a cross in the column “matrix” and the line “change” in the legend to the individual figures of Fig. 9.
3. Mutation type 3 (“replace replicator”): This procedure becomes active only when a replicator dies out. It replaces it with the randomly chosen copy of a survivor and replaces one of the also randomly chosen positions of its interaction vector with a random number z (0<z≤1). The initial frequency is adjusted as described. When this procedure is used, there is a cross in the “replicator” column and the “replace” row in the legend to the individual figures of Fig. 9.
4. Mutation type 4 (“change replicator”): This mutation type also becomes active when a replicator dies out. When a randomly selected copy of a survivor takes the place of the extinct one, a randomly selected position of its interaction vector is mutated in the way already described. As already explained in point 1, the initial frequency is changed. If this procedure is used, there is a cross in the column “replicator” and the line “change” in the legend to the individual figures of fig. 9.

The difference between mutation type 1&2 on the one hand and 3&4 on the other hand is that with 1&2 also an extinct replicator can be “revived” by replacement or mutation (of course an actually unrealistic, but just possible, procedure), or a survivor is changed, whereby the original disappears. In 3&4, on the other hand, a survivor is changed, but the original remains.

The four basal mutation types can be combined in various ways, e.g. by alternating them or by using them with different probabilities. For Fig. 9, the resulting different evolutionary processes were each repeated 100 times (while 10000 repetitions were performed for the study of selection). Each blue dot in Fig. 9 corresponds to the result of evolution, each yellow one to that of selection only. One yellow and one blue dot each form a pair, but this cannot be seen in this kind of representation. In Fig. 9 only the result of those evolutionary procedures is shown, where the distributions of “selection only” and “evolution” are clearly different, i.e. which have caused something. The single mutation types were carried out, as well as all combinations of two of them, and finally the combination of all four. Procedures that only change the matrix without making a copy of a survivor (mutation type1 and 2) were not successful if not combined with others. One algorithm increases synreplicator complexity without increasing mean fitness (change replicator). Another primarily changes the mean fitness (change matrix, change replicator), and some increase both. The lower right image shows the result of an algorithm not yet described, where the new replicator combines the interaction vectors of two survivors. As can be seen, this also yields success.

The same evolutionary algorithms were also applied to recombinator systems (not shown). Here, however, none of the mutation types and no combination of them brought even the slightest success, i.e. an increase in mean fitness and/or synreplicator complexity was not achieved.

In this work, the concept of entropy has been extended to the case of frequency-dependent selection and it has been shown for recombinator systems that the total entropy change can be calculated by adding the contributions of the individual gene loci. However, the question of how this extended concept of entropy can be incorporated into information theory has remained open. For replicator and recombinator systems with randomly selected interaction matrix values, it has been shown how the number of replicators surviving on average, the mean fitness, and the entropy change depend on the number of replicators (recombinators) n initially present, on their initial frequencies x_i_, and on certain matrix properties (symmetry). It has been shown that the survival of several to many replicators or recombinators is the norm rather than the exception. In a paper on evolution and cooperation Nowak^16^ writes: “Natural selection implies competition and therefore opposes cooperation unless a specific mechanism is at work”. This view is not confirmed by the present results. It seems to be rather a consequence of a teleological view. Of course, one can criticize the approach of using random interaction matrix values. Catalytic abilities are much more widespread in molecules than autocatalytic ones, so that, at least in prebiotic evolution, it seems justified not to assume that a_rs_ must be larger when r=s than when r≠s.

Special cases were studied for entropy, but their systematic classification was left to the future. Finally, how evolution changes such systems has also been analyzed, showing that different evolutionary algorithms in replicator systems can lead to an increase in mean fitness, or to an increase in the number of survivors, or both. This also rather contradicts the classical view, which Nowak^16^ formulates as follows: “Evolution is based on a fierce competition between individuals and should therefore reward only selfish behavior”. This view has haunted us for more than 150 years now and has given us Spencer’s Social Darwinism, Galton’s Eugenics, and Haeckel’s race theories of Aryan superiority. In fact, however, it represents an artifact that arises when one can analyze only very simple replicator systems, as was the case for Darwin’s and even Fisher’s contemporaries, and generally before the invention of the computer. In more complex replicator systems, cooperation follows naturally as a consequence of selection and evolution. Evolution is not purposeful, it does not reward anything, not even selfish behavior. Replicators also do not “want” to survive, even if they can program their carriers to develop a survival instinct. Evolution should no longer be seen in this way, but as it actually is: merely as a process of acquiring information.

All the evolutionary algorithms described fail in recombinator systems if at the beginning the interaction matrix values are randomly chosen. This does not mean, of course, that recombinators cannot evolve, for eukaryotes are highly successful representatives of terrestrial evolutionary events despite, or rather because of, sexual reproduction. The clarification of what alternative evolutionary algorithms come into play in the context of sexual reproduction and what conditions must be met for it to be successful is also left to future work.

## Supplement 1: Algorithms for the determination of the allele index k, if the genotype index i and the gene loci index j are known

The value of the allele index k (k=1…l) depends on the genotype index i (i=1…n) and gene loci index j (j=1…m). The number of genotypes n is given by

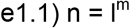

where l is the number of alleles per gene locus and m is the number of gene loci. We assume without restriction of generality that l is the same for all gene loci (allele frequency can be zero, of course).

The problem of getting from the genotype index, which we write as a decimal number, to the gene locus and allele index corresponds to the algorithm that allows the transformation from the decimal system to a number system with base l. To do this, the genotype index is divided continuously (with integer result) until the quotient is zero^18^. The rest of each division, which is between 0 and l-1, forms the digits of the resulting number. In the following example, let i=20, l=2, and m=5. From l and m, it follows according to Eq. e1.1 that there are then 32 different genotypes. Since our index i does not start at zero but at one (as is common in mathematics in contrast to computer science), we have to calculate with i-1, i.e. with 19:

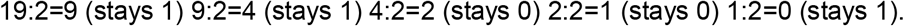

Also our allele index does not start at zero and therefore by adding 1 for the allele indices of the gene loci 1 to 5 we finally get: k=2, 2, 1, 1, 2 for the genotype index 20 (note: for the representation of 19, i.e. the twentieth number as a dual number, the order is reversed, furthermore with 0 instead of 1 and 1 instead of 2: 10011).

If one does not want to determine k for all gene loci, just to know the allelic index for a specific j, one can use e.g. the “k-wheel” (Fig. F1.1).

**Fig. f1.1:**
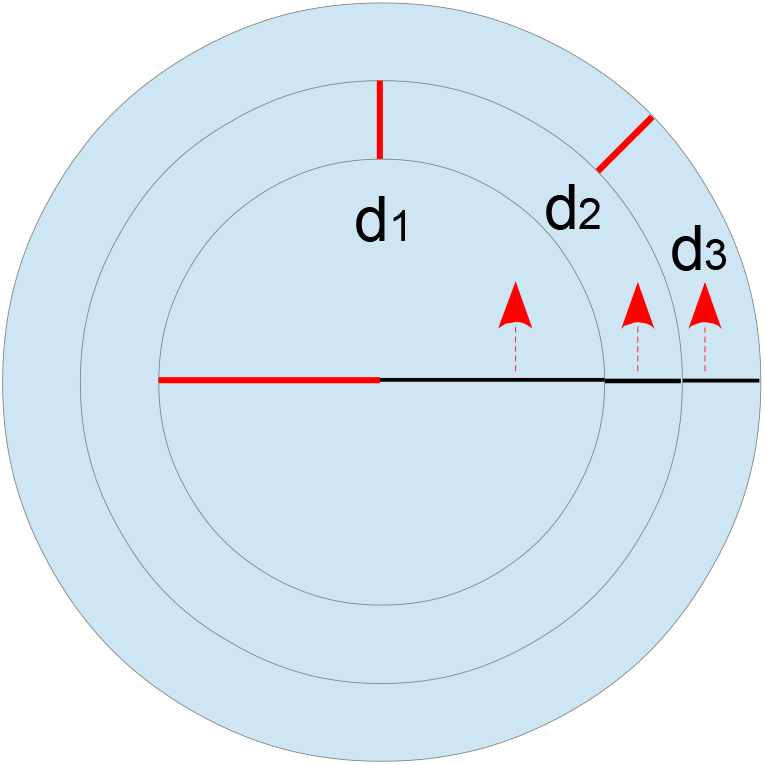
The k-wheel for finding k given i and j. In this example, l=2 and j=1 (inside) to j=3 (outside).

In principle, the wheel is rotated around a circle part d_j_, which is given by:

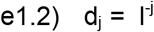

If there are two alleles per gene locus, e.g. d_1_=1/2, d_2_=1/4, d_3_=1/8 and so on. Now turn the wheel i-1 times around the arc d_j_, of course always in the same direction. In doing so, we must take into account the cyclic nature of our algorithm. The easiest way to do this is to remember that every real number z can be viewed as the sum of an integer ^G^[z] and a residue ^g^[z]:

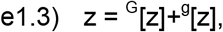

By considering only the residue ^g^[z], we ignore how many times the wheel has already made a full rotation (we could also use a trigonometric function for this, but it’s easier this way). ^g^[z] is the same for z=1.3, z=2.3, and z=5.3, for example, namely 0.3. We now define f_ij_ as follows:

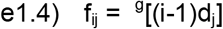

and finally obtain k as the integer part of the product of l and f_ij_, added by one:

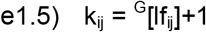

The following table shows the values for the index k_ij_ given i and j (maximum three gene loci, i.e., m=3), if there are two alleles at each gene locus (l=2; thus, there are two possible values for k, namely, 1 and 2).

**Table t1.1:**
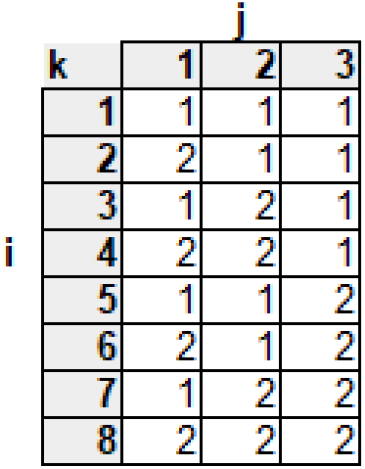
The allele combinations of the i-th genotype X_i_. For example, X_4_ corresponds to the B_jk_ combination: B_12_B_22_B_31_.

In the practical implementation of this algorithm, however, one may encounter the problem that the computer stores e.g. 1/3 as 0.3333… and therefore 3(1/3) as 0.9999…. When determining the integer part, the responsible function may therefore return the value 0 instead of 1, as would be correct. One can solve this problem by adding a very small number to the calculation of f_ij_, e.g. f_ij_ = ^g^[(i-1)d_j_ + 0.000001]. If one intends to calculate all possible values anyway, it is better to choose another algorithm, such as one based on a recursive function.

## Supplement 2: Addability of the mean fitness of the gene loci if the total fitness is addable. Example with two gene loci to two alleles

For the mean fitness ω holds:

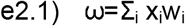

or in the case of 4 geno(pheno)types

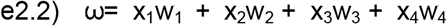

We assume that the four genotypes are the possible combinations of two alleles at each of two gene loci. After recombination has occurred, the relative frequency of X_i_ is given by x_i_. Of course, Σ_i_x_i_=1 and Σ_k_b_jk_=1 hold for all j (b represents the frequency of allele B):

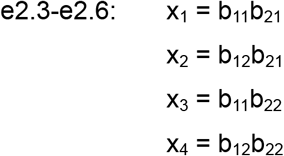

With additive fitness, w_i_ of genotype X_i_ is calculated as follows:

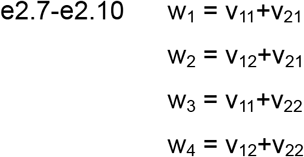

Where v_jk_ is the fitness of the kth allele of the jth gene locus. The frequency of alleles B_jk_ is given by (where Σ_k_b_jk_=1 for each gene locus j):

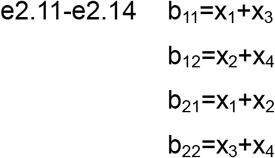

as follows from e2.3-e2.6. We now assume that in case of additive fitness also the mean fitness ω behaves additively, i. e.:

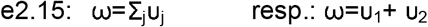

with

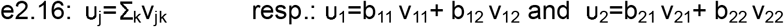

This is what we have to show. We start according to eq. e2.15 and e2.16 with:

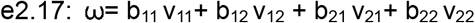

We now replace allele frequencies with genotype frequencies according to e2.11-e2.14:

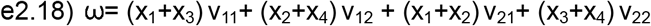

from which follows after a short calculation:

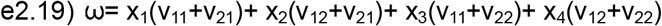

Now we can make a second substitution after e2.7-e2.10 and get:

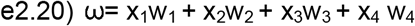

which corresponds to eq. e2.2, showing for a simple example that the mean fitness is (unsurprisingly) additive when the fitness is additive. In all other cases, however, this does not hold, while Eq. 7 always holds regardless of the selection value landscape! By the way, Fisher^4^ was able to show in 1930 that in the case of additive fitness, the mean fitness always increases by selection until it reaches the maximum. Multi-peaked selection value landscapes do not occur in this case, so that selection cannot catch itself in an intermediate optimum.

